# Mutual Annotation-Based Prediction of Protein Domain Functions with Domain2GO

**DOI:** 10.1101/2022.11.03.514980

**Authors:** Erva Ulusoy, Tunca Doğan

## Abstract

**Motivation:** Identifying unknown functional properties of proteins is an important task for understanding their roles in both health and disease states. The domain composition of a protein can reveal critical information in this context, as domains are structural and functional units that dictate how the protein should act at the molecular level. The expensive and time-consuming nature of wet-lab experimental approaches prompted researchers to develop computational strategies for predicting biomolecular functions. Biological ontologies, such as the Gene Ontology (GO), which provide a standardized vocabulary of information about biological entities, are frequently employed in protein function prediction.

**Results:** In this study, we proposed a new method called Domain2GO that predicts associations between protein domains and GO terms, thus redefining the problem as domain function prediction, using documented protein-level GO annotations together with proteins’ domain content. To obtain reliable associations, co-annotation patterns of domains and GO terms in the same proteins are examined using statistical resampling. An ablation study was conducted to compare the predictive performance of various implementations of Domain2GO, differing from each other by the utilized statistical measure (e.g., information theory inspired similarity measures and the ones calculated by the expectation-maximization algorithm). As a use-case study, examples selected from the finalized domain-GO term mappings were evaluated for their biological relevance via a literature review. Then, we applied the proposed method to predict presently unknown protein functions by propagating domain-associated GO terms to proteins annotated with these domains. For protein function prediction performance evaluation and comparison against other methods, we employed Critical Assessment of Function Annotation 3 (CAFA3) challenge datasets. The results demonstrated the high potential of Domain2GO, particularly for predicting molecular function and biological process terms, along with advantages such as producing interpretable results and having exceptionally low computational costs. The approach presented here can be extended to other ontologies and biological entities in order to investigate unknown relationships in complex and large-scale biological data.

**Availability and implementation:** The source code, datasets, results, and user instructions for Domain2GO are available at https://github.com/HUBioDataLab/Domain2GO.

## 1. Introduction

To comprehend biological mechanisms and develop effective treatment strategies for diseases, it is necessary to have knowledge of both the molecular functions of proteins and their roles in large-scale biological processes. Experimental function identification techniques cannot keep up with the increasing amount of sequence data due to their expensive and time-consuming nature. The ever-expanding output of high throughput sequencing is stored in public databases such as the UniProt Knowledgebase (UniProtKB) (UniProt Consortium, 2020), where the vast majority of protein sequences (%99, as of September 2022) are missing experimental and/or annotation.

Considering that the biological functions of proteins are multifaceted and there is a vast amount of scientific knowledge on this topic, it is important to define this knowledge in a systematic and machine-readable way for standardization. There are various classification systems that have been created to meet this need. The Gene Ontology (Ashburner et al., 2000; Carbon et al., 2020) is the most extensive and widely used resource, providing a standardized vocabulary of information on the functions of gene products. The Gene Ontology defines different aspects of protein function in three sub-ontologies (i.e., molecular function, biological process, and cellular component), which contain a total of 43,791 terms (as of November 2021). GO terms of each sub-ontology are hierarchically linked to each other with well-defined relationships, forming a directed acyclic graph (DAG).

Protein family and domain databases such as InterPro (Hunter et al., 2009) stand as comprehensive resources of protein signatures, which can be used to infer the structure and function of unknown proteins. InterPro2GO mappings (Camon et al., 2005), created by the InterPro team, aim to provide high-quality curated functional annotations directly for domains. Even though this approach has been used since 2002, the annotation coverage of InterPro v83.0 only remains at 41% (15,839 out of 38,345 InterPro entries are associated with at least one GO term).

A variety of computational approaches have been developed for predicting protein functions over the years (Unsal et al., 2022; Rifaioglu et al., 2019, 2018; Doğan, 2018; Kulmanov et al., 2020; Zhao et al., 2022), to serve the ultimate aim of attaining a complete understanding of biological mechanisms. The idea behind these approaches is that annotations can be transferred among proteins sharing similar characteristics (e.g., sequence, structure, protein-protein interactions, expression profiles, etc.). One efficient class of methods utilizes the domain-based approach to address the challenging task of protein function prediction. Domain composition of a protein is one of the key indicators of its function(s). There are earlier studies that aim to investigate similar domain compositions in different proteins in order to infer their functional properties. Forslund & Sonnhammer (2008) implemented this idea as a simple and intuitive tool that makes use of available domain architecture data to transfer annotations between distant proteins. While most of these methods consider only the domain content of proteins (Das et al., 2015; Rojano et al., 2022), there are studies that involve attributes such as order, position, and recurrence of domains to measure similarity between domain architectures (Doğan et al., 2016). Integrating multiple sequence-related properties, including the domain architecture, using machine learning algorithms has been shown to improve the function prediction performance (You et al., 2018, Kulmanov et al., 2020).

The Critical Assessment of Function Annotation (CAFA) (Radivojac et al., 2013; Jiang et al., 2016; Zhou et al., 2019) is a time-based challenge designed to meet the need for assessing the performance of protein function prediction approaches in a comprehensive and objective manner. At the beginning of each round of the CAFA challenge, organizers provide a target set that contains proteins whose functions are unknown. Participants predict the functions of proteins in this dataset and submit them. During a data collection period, target proteins that gain experimental annotation accumulate a benchmark set, which is then used to evaluate the performance of participating methods. The CAFA challenge has been widely adopted in the protein science community and has become a community-driven worldwide benchmark effort with three completed and one ongoing installment as of September 2022.

Although each of these function prediction methods helps to bridge the gap between sequencing and function annotation capabilities, protein function prediction remains an open problem, as evidenced by recent CAFA experiments. A limitation of the machine learning-based models, which are among the best-performers in CAFA challenges, is that it is hard to comprehend the underlying decision mechanisms behind predictions. Therefore, most of these models are problematic in terms of interpretability. Another limitation is related to computational complexity. Although hybrid models that combine multiple data types and/or algorithms could achieve high prediction performance, they come at a high computational cost, which makes them incapable of scaling up to large-scale datasets such as the entire UniProtKB.

To address the issues outlined above, we present a new method, Domain2GO, that utilizes the associations between protein domains and functions in order to infer GO term annotations for uncharacterized proteins (Fig. 1). Domain2GO links two types of relational information to each other: domain-protein and protein-function, in order to obtain GO term predictions for both InterPro domains and full proteins. InterPro domains and GO terms that display statistically significant co-occurrence patterns on different proteins are found to be related and included in our association list called “Domain2GO mappings”. Accordingly, the GO terms of interest are used to annotate the proteins that contain the corresponding domains, resulting in protein function predictions. We evaluated the results of our approach by comparing the obtained predictions with the manually curated domain-function mappings from InterPro2GO (Camon et al., 2005). Additionally, we employed Domain2GO to predict the functions of the target proteins from the third version of the CAFA challenge (CAFA3) and compared the prediction performance against the baseline methods in order to examine the capability of Domain2GO to produce large-scale protein annotations as a downstream secondary task. As a use-case study, we examined selected Domain2GO pairs in order to investigate whether the results have biological relevance. The finalized Domain2GO mappings were used to obtain function predictions for the proteins in the whole UniProtKB/Swiss-Prot database.

**Figure 1.**
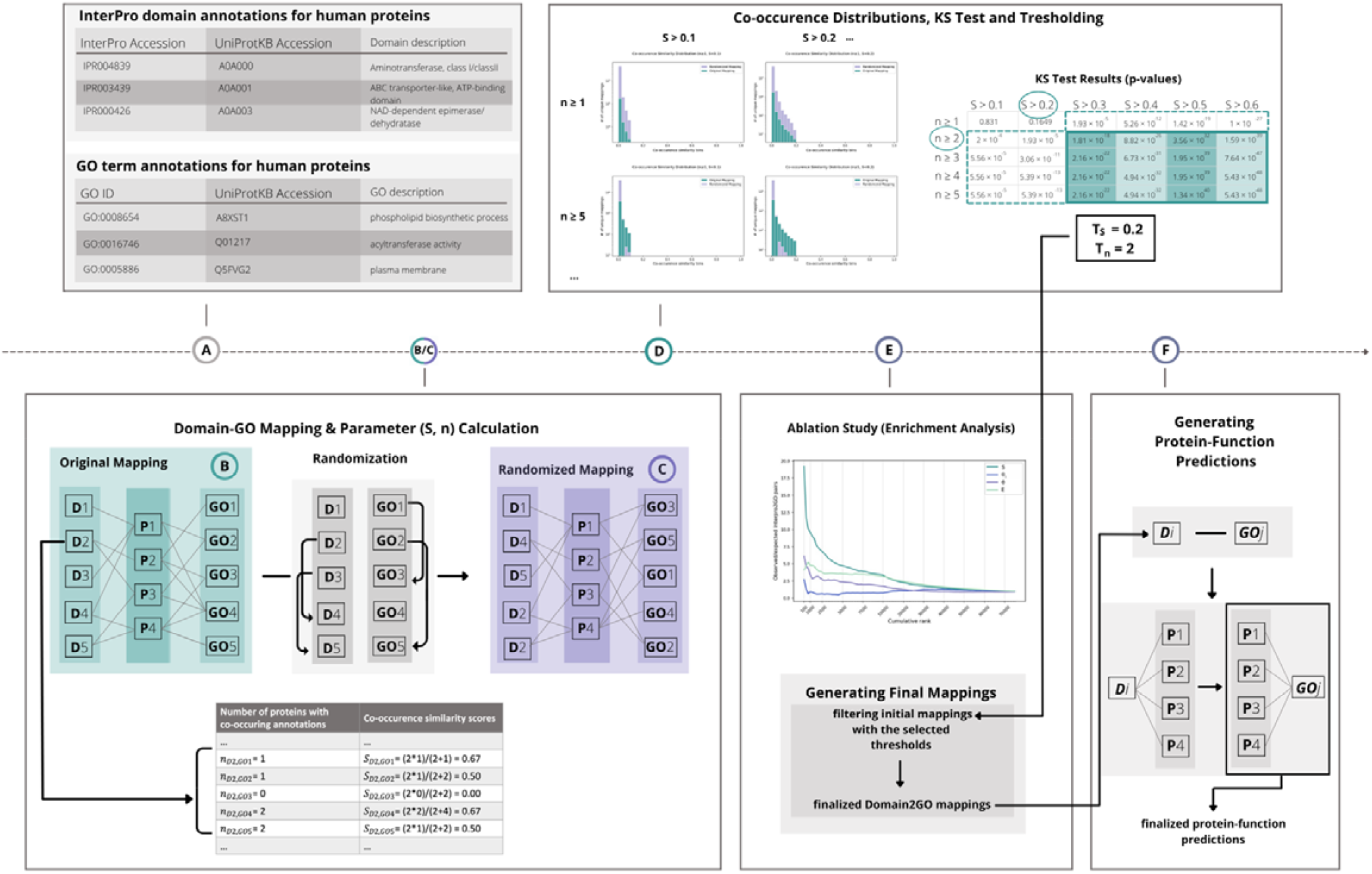
Schematic representation of the proposed method. **(A)** The source datasets were downloaded and organized; **(B)** initial mapping between the InterPro domains and GO terms were obtained and the mapping parameters were calculated; **(C)** generation of the randomized annotation and mapping sets were constructed; **(D)** co-occurrence similarity distributions were plotted, and thresholds were selected based on statistical resampling; **(E)** an ablation study was conducted by calculating the enrichment of top predictions ranked by different statistical measures, and finalized Domain2GO mappings were generated by filtering initial mappings; **(F)** protein function predictions were generated by propagating Domain2GO mappings to target proteins.

## 2. Methods

The overview of the proposed methodology is shown in Fig. 1. First, we prepared the domain and GO annotation datasets and used the UniProt accessions in both sets to map InterPro domains to corresponding GO terms (Fig. 1A and 1B). We identified each pair as a(n) initial/raw Domain2GO mapping. The training part of the system consists of two statistical score calculation steps. First, we calculated a co-occurrence similarity measure (S) for each Domain2GO mapping. This measure was utilized in statistical resampling (via the Kolmogorov–Smirnov test), which is employed to differentiate significant mappings from the ones that occurred by chance (Fig. 1C and 1D). Second, we applied the expectation maximization (EM) algorithm (Dempster et al., 1977), which is a method for obtaining a maximum-likelihood estimate of the probability of association (θ), and the confidence of its probability (E), independently for each Domain2GO mapping. All Domain2GO mappings were ranked independently by four of the calculated measures (i.e., S, θi, θ and E) and the enrichment of manually curated Domain2GO annotations in the highest-ranking mappings was analyzed. The initial mappings were filtered using the thresholds established in previous steps, and the resulting mappings were employed to generate protein function predictions (Fig. 1E).

### 2.1. Dataset

In order to generate domain-function mappings, we combined domain-protein and protein-function associations. For the domain-protein annotation dataset, the UniProtKB-InterPro entry annotation file was downloaded from the InterPro database version 85.0. This file contained 615,060,473 annotations between 38,913 InterPro entries of all types and 179,520,976 UniProtKB proteins. After eliminating all InterPro entry types other than domains, the final version of the file contained 222,753,705 annotations between 11,233 InterPro domain entries and 121,492,444 UniProtKB proteins.

The second dataset contained protein function annotations, which were obtained by following a procedure utilized in (Doğan et al., 2016), as follows: GO annotations with manual and experimental evidence codes (i.e., all except electronic annotations -IEA, due to their questionable reliability) were downloaded from the UniProt-GOA database version 193 (Huntley et al., 2015) for the protein entries in the UniProtKB/Swiss-Prot database (UniProt Consortium, 2021). The GO terms in the dataset were propagated to their parent terms based on their “is a” and “part of” relationships, to enrich the annotations. In its final form, the dataset had 6,998,018 annotations between 125,309 proteins and 30,959 GO terms.

### 2.2. Calculating the co-occurrence similarities

We used the UniProt IDs as the key to connect two datasets and generate domain-GO term mappings. If a UniProt ID is shared between (co-annotated with) an individual domain and a GO term, we identify this pair as a(n) initial/raw Domain2GO mapping. This way, 2,069,796 mappings were obtained between 8,642 domains and 28,420 GO IDs. In the first step of the training procedure, a filtering operation was performed to eliminate mappings resulting from the coincidental occurrence of a domain and GO ID on the same protein. For this, we calculated a co-occurrence similarity score (Eq. 1.), inspired by an information-theoretic definition of similarity (Lin, 1998) for each domain-GO mapping.

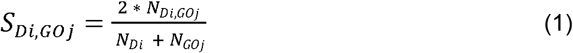

where S_Di, GOj_ is the co-occurrence similarity of a domain i and GO term j pair, N_Di, GOj_ is the number of proteins where i and j are annotated together, N_Di_ is the total number of proteins with the annotation “i” and N_GOj_ is the total number of proteins with the annotation “j”.

An example of the co-occurrence similarity calculation is shown in Fig. 1B and C. A high degree of co-occurrence similarity indicates that the number of proteins to which a protein domain and GO terms are co-annotated is significantly greater than the total number of proteins to which these terms are individually annotated. Therefore, a threshold co-occurrence similarity value can be set to differentiate the relevant Domain2GO pairs from the ones that occurred by chance. Assuming that some pairs with a high similarity score can still be random, especially when N_Di_ and N_GOj_ are very low, we employed n (N_Di, GOj_); the number of proteins with co-occurring i and j annotations, as a second parameter. Accordingly, Domain2GO pairs with both high S and n values are considered to be more reliable.

In order to assign optimal S and n values to be used as thresholds, we compared the co-occurrence similarity distribution obtained from Domain2GO mappings to the score distribution of randomly generated domain-GO pairs. The idea behind comparing these distributions is that the frequency of mappings with high and low S values should differ between the original and random mapping sets. For this, a randomized mapping table was created by shuffling domain and GO term columns on the original Domain2GO mappings (Fig. 1C), then the S and n score calculation steps were repeated for the newly generated random Domain2GO mappings.

30 candidate threshold value combinations were generated for S (i.e., S < 0.1, < 0.2, < 0.3, < 0.4, < 0.5, and < 0.6) and n (n>= 1, >= 2, >= 3, >= 4, and >= 5). The co-occurrence similarity distribution histograms of the mappings with S and n values higher than the candidate thresholds were plotted for both the original and randomized mapping sets. As a last step, we employed a widely used non-parametric test, Kolmogorov–Smirnov (KS) (Lilliefors, 1967; Hollander, Wolfe & Chicken, 2013), which compares two samples and estimates whether they are drawn from the same distribution by quantifying the distance between the samples. The null hypothesis states that the samples are drawn from the same distribution. The KS-test was applied to 30 different co-occurrence similarity distributions, which were plotted for each of the candidate threshold value combinations. The minimum value pair that is able to show that the original and random mapping sets are drawn from significantly different distributions (i.e., the lowest values the null hypothesis is rejected at) was selected as the official thresholds.

### 2.3. Expectation maximization (EM)

#### 2.3.1. Estimating association probabilities of Domain2GO pairs

For the second step of the training process, we applied the EM algorithm (Dempster et al., 1977) which was implemented in earlier studies for different prediction purposes, such as domain-domain interaction prediction (Riley et al., 2005; Deng et al., 2002).

The algorithm was initialized by obtaining an estimate of the association probability for each Domain2GO mapping. In order to do this, all the mappings were represented in a data structure, *C*, which was initialized by setting all C_p_ ^Di, GOj^ = 1 (Eq. 2), assuming that all possible domain-GO term pairs that are co-occurred on the same protein can be associated with each other. Based on *C*, two other statistics were defined. M_Di, GOj_ (Eq. 3) is the number of proteins where GO term j is co-annotated with domain i. N_Di, GOj_ (Eq. 4) is the number of proteins where j is not co-annotated with i. Finally, Z_Di, GOj_ is defined, which is the number of unique proteins where j is annotated to, but i is not annotated to, and vice versa.

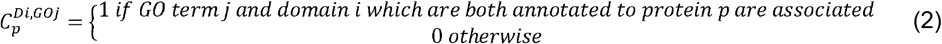

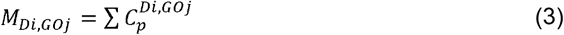

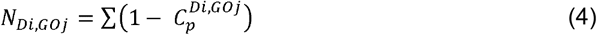

M_Di, GOj_ was initialized with the number of proteins where i and j are annotated together and N_Di, GOj_ was initialized with zero based on their equations, as all initial C_p_^Di, GOj^= 1. As EM proceeds, M and N change based on modified values of C, yet Z is a fixed value.

After these measures are defined, we calculated the initial estimate of the θ_Di, GOj_ (i.e., the probability that GO term j and domain i is associated) (Eq. 5)

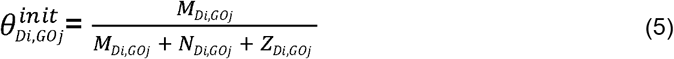

Initialized values of θ and C were then used to estimate the likelihood L (Eq. 6) of the Domain2GO mappings.

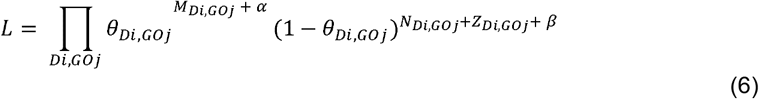

Where α and β are arbitrary pseudocounts to ensure that obtaining too high or too low θ_Di, Goj_ values were not due to the low frequencies of i and j, but to the large numbers of co-occurrences indicating the potential association of them. After initialization, we followed the four basic steps of the EM algorithm:

The expected value of C_p_ ^Di, GOj^ was calculated for each Domain2GO co-occurrence:

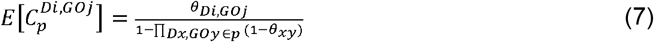

1. M_Di, GOj_ and N_Di, GOj_ were calculated for each Domain2GO pair from the expected value of C_p_ ^Di, GOj^.
2. θ_Di, GOj_ was re-estimated from the expected values of M_Di, GOj_ and N_Di, GOj_.
3. Likelihood (Eq. 6) was re-estimated.

We repeated these steps until the likelihood (Eq. 6) stops increasing noticeably. The probability matrix of all Domain2GO domains, θ, was obtained as the result of the EM algorithm. A high θ_Di, GOj_ value shows that the mapped domain and GO term have a high probability of being associated. At this point, the issue mentioned in Section 2.2 emerges again, due to the possibility of observing pairs with high θ just by chance when some domains and GO terms were annotated to a high number of proteins. Furthermore, a domain-GO mapping, in which the corresponding GO term is highly specific, may have a low θ just because the domain and GO term are both annotated to a low number of proteins. A likelihood ratio test was performed in order to differentiate highly significant domain-GO mappings from the ones that could be observed just by chance.

#### 2.3.2. Computing E scores (Likelihood Ratio Test)

A new probability matrix, 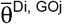,was defined by running the EM algorithm again, under the model that all instances of i-j pair given a θ_Di, GOj_ of zero. This resulted in a maximum likelihood estimate of all Domain2GO mappings excluding D_i_ -GO_j_ pair. A measure of evidence, E_Di, GOj_, is calculated by using the 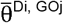 matrix defined specifically for each pair (Eq. 8).

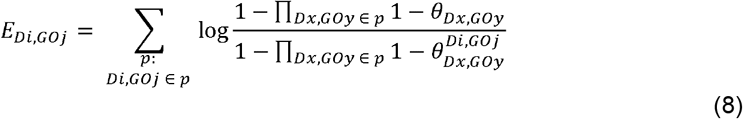

Where θ is the probability matrix of all domain-GO mappings and 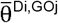 is the probability matrix of all domain-GO mappings under the model that the probability of given i-j mapping (θ_Di, GOj_) is equal to zero.

While θ can be considered as the probability of a mapping, the E score aims to detect highly specific Domain2GO mappings with extensive evidence in the co-annotation data. The number of instances of a mapping should increase the confidence in that annotation, therefore, a high E_Di, GOj_ is expected for a pair with high θ. More importantly, a pair with low θ can also have a high E_Di, GOj_ if θ is relatively high in comparison with the competing 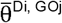 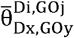. On the contrary, a low E_Di, GOj_ implies that this pair is not any better at explaining co-annotations than competing hypotheses (other potential pairs which are co-annotated to the same protein).

### 2.4. Competing function prediction methods

Two baseline methods, Naive classifier and BLAST, were built in order to compete with the co-occurrence similarity and E score-based function predictions of Domain2GO. Both of the baseline methods were constructed as explained in the CAFA experiments (Zhou et al., 2019), using the Swiss-Prot annotations that existed before the CAFA3 annotation collection period (date: 2016/09). In addition, we include InterPro2GO annotations as another competing model that considers domain-based knowledge. It was possible that the current InterPro2GO release includes the annotations obtained within the CAFA3 benchmark collection period. In order to avoid giving InterPro2GO an unfair advantage, we used the September 2016 InterPro2GO release. Domain-GO term pairs from the InterPro2GO data were employed to generate protein-function predictions on the CAFA3 target protein set. As InterPro2GO annotations are created through manual curation, all prediction scores were set to 1.

It is important to note that only annotations produced during the annotation collection period are used to generate the CAFA benchmark set. Hence, a prediction may not be included in the CAFA benchmark dataset even though it is verified after the end of the challenge. These predictions are evaluated as false positives during evaluation.

### 2.5. Performance evaluation metrics

We employed the CAFA3 challenge (Zhou et al., 2019) benchmark set to investigate whether Domain2GO can be extended to predict protein functions, even though it is primarily proposed with the aim of predicting functions of domains.

The Domain2GO mappings were recalculated using data from September 2016, to comply with the calendar of the CAFA3 experiment and thus yield a fair comparison with other methods. The mappings were then used to generate function predictions for the CAFA3 target set proteins. Here, we used the protein-centric evaluation mode, which measures the accuracy of assigning GO terms to a protein by calculating two basic evaluation metrics: Fmax and minimum semantic distance (Smin) (Eq. 9 and 12).

Fmax (Eq. 9) corresponds to the maximum of the F-scores that are computed using precision - pr (Eq. 10) and recall - rc (Eq. 11) values at each prediction score threshold (τ).

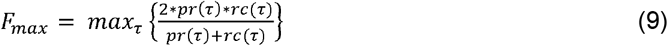

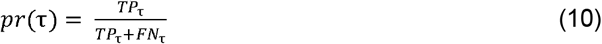

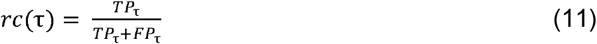

Where *TP*τ, *FN*τ and *FP*τ represent the number of true positives, false negatives and false positives at the prediction score threshold τ.

Smin corresponds to the minimum semantic distance between real and predicted annotations computed using remaining uncertainty (ru) and misinformation (mi) values over all prediction score thresholds. Smin (Eq. 12) evaluates the predictive performance by taking into account the information content of GO terms, down-weighing the less informative annotations as they are not as valuable.

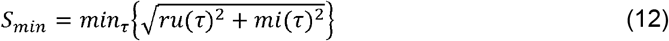

Detailed information regarding the definition and calculation of these metrics can be found in (Jiang et al., 2016). The prediction score thresholds are selected by the CAFA3 evaluation protocol to generate binary predictions from submitted continuous prediction scores. Additionally, we report a secondary metric, coverage, corresponding to the percentage of the proteins in the benchmark that are predicted by the proposed method.

Co-occurrence similarity and E score results were evaluated as two distinct models to determine their predictive capabilities. To calculate the performance using the CAFA3 protocol, prediction scores must be within the range of (0.00,1.00]. For the first prediction score set, we directly used the co-occurrence similarity scores (which were already set to the required range) calculated in Eq. (1). If a protein with an existing domain i annotation receives the GO term j prediction, then this protein-function pair obtains the prediction score S_Di, GOj_. For the second prediction score set, we used the E values calculated in Eq. (8), after normalization by min-max scaling. Similar to the first one, if a protein with domain i annotation receives the GO term j prediction, then this protein-function pair receives the normalized E_Di, GOj_ value as its prediction score.

## 3. Results and Discussion

### 3.1. Statistical analysis of the mappings

The original and randomized Domain2GO mapping generation process is explained in detail in Section 2.2. The procedure resulted in 2,069,796 unique initial/raw mappings for the original and 1,927,988 unique mappings for the randomized set, between 8,642 domains and 28,420 GO terms.

Table 1 displays the numbers of Domain2GO pairs, domains, and GO terms in the original and randomized mapping sets at different co-occurrence similarity thresholds. It is observed that the original and randomized mapping sets have a similar number of mappings when the S threshold is lower. However, the gap between them widens when we look up to higher S thresholds. This is expected given that a domain i and a GO term j are much less likely to co-occur on the same protein when their indices are randomized. This also explains the lower number of domains and GO terms at higher S values for the randomized mapping set.

**Table 1.** Statistics of the initial original and randomized Domain2GO mappings (n≥1 for all values).

| Threshold (S) | # of mappings |  | # of unique domains |  | # of unique GO terms |  |
| --- | --- | --- | --- | --- | --- | --- |
|  | Original mapping | Random mapping | Original mapping | Random mapping | Original mapping | Random mapping |
| = 1 | 470 | 35 | 319 | 35 | 350 | 34 |
| $\geq 0.9$ | 902 | 35 | 574 | 35 | 679 | 34 |
| $\geq 0.8$ | 1,667 | 35 | 997 | 35 | 1,211 | 34 |
| $\geq 0.7$ | 2,511 | 35 | 1,359 | 35 | 1,752 | 34 |
| $\geq 0.6$ | 4,278 | 212 | 2,076 | 207 | 2,847 | 195 |
| $\geq 0.5$ | 6,837 | 570 | 2,917 | 542 | 4,398 | 479 |
| $\geq 0.4$ | 10,619 | 1,096 | 3,741 | 1,016 | 6,392 | 832 |
| $\geq 0.3$ | 16,801 | 1,811 | 4,602 | 1,622 | 9,122 | 1,240 |
| $\geq 0.2$ | 32,028 | 6,626 | 5,775 | 2,986 | 14,236 | 4,929 |
| $\geq 0.1$ | 78,477 | 30,459 | 6,870 | 5,509 | 21,466 | 12,692 |
| $\geq 0.0$ | 2,069,796 | 4,584,047 | 8,642 | 8,642 | 28,420 | 28,420 |

Co-occurrence similarity distributions for the original mapping set are shown in Fig. 2 as histograms for selected n values. It has been observed that mappings with low co-occurrence similarity, which have a high probability of being paired by chance, are accumulated in the distributions of low n values (e.g., 1 and 5). When higher n thresholds were applied, the number of these low-confidence mappings was drastically reduced, indicating that the reliability of the mappings in the dataset has increased. 99.98% and 99.99% of the mappings with a co-occurrence score lower than 0.0225 (i.e., the first three bins of the histograms) were eliminated at n≥25 and n≥75 respectively (Fig. 2c and 2d). Regardless of this improvement, it is important to note that when extreme n thresholds (i.e., 25 and 75) were applied, only a small portion of the initial mapping set remained. Moreover, it is possible to observe mappings with extremely low S values even at these stringent n threshold values. These observations demonstrate the necessity of combining S and n values to filter out unreliable mappings.

**Figure 2.**
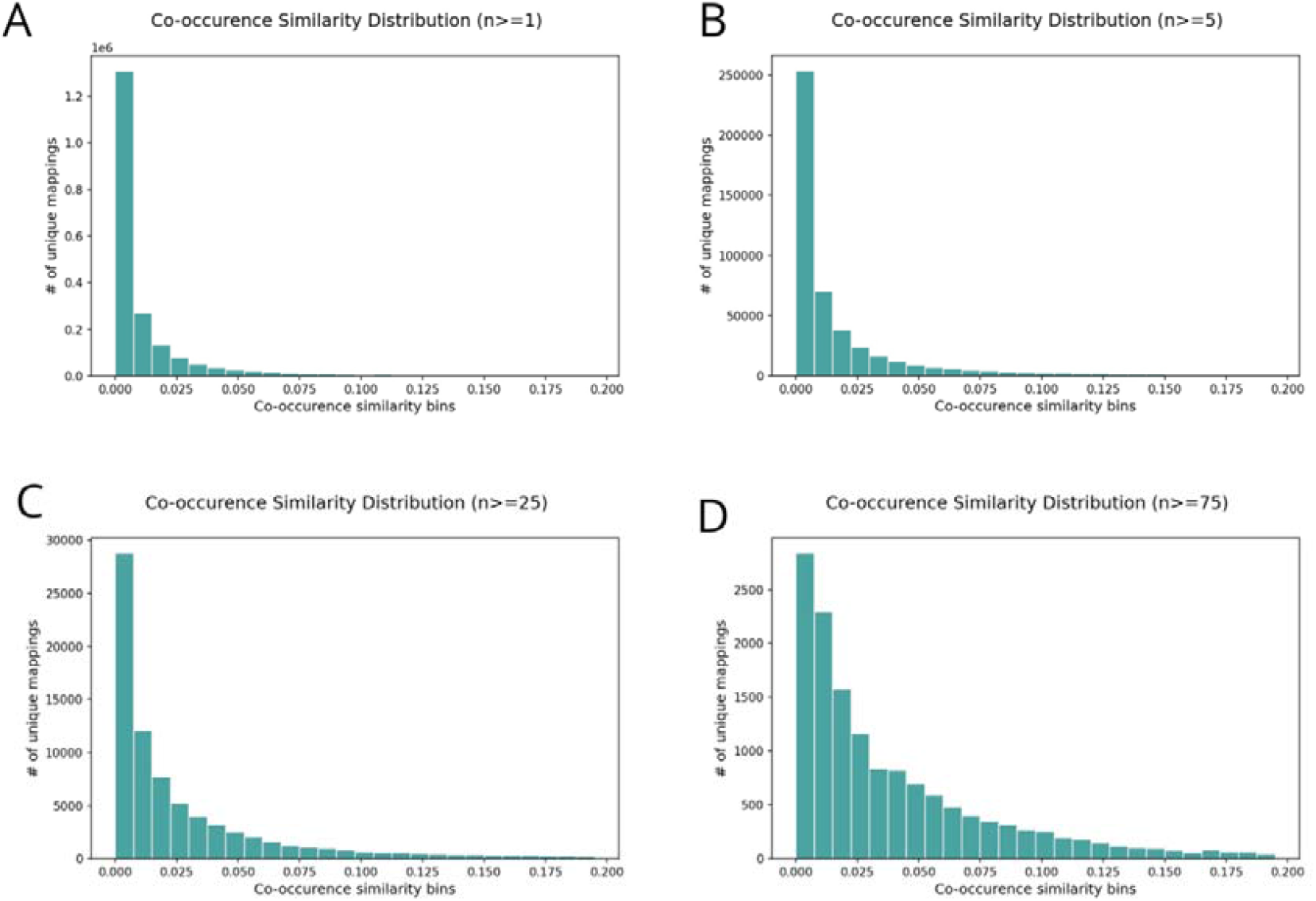
The co-occurrence similarity distributions of the initial/raw original Domain2GO mappings. Each plot is drawn for a different threshold of the number of co-annotated proteins (n); **(A)** n≥1, **(B)** n≥5, **(C)** n≥25 **(D)** n≥75.

We utilized the KS test to determine the optimal S and n threshold pair for distinguishing relevant mappings from the remainder. By combining arbitrarily selected n and S values, 30 co-occurrence distributions were generated for both the original and randomized mapping sets (Table 2). The KS test was applied to the remaining data points from both distributions following the filtering operation by applying the corresponding S and n threshold pair. The minimum value pair that rejects the null hypothesis, which states that the two samples (i.e., the original and the randomized mapping set) are drawn from the same distribution, was selected to be used as finalized thresholds. As can be seen in Table 2, the minimum value pair that yielded the required statistical significance (i.e., p-value < 0.01) is n ≥ 2 and S < 0.1. The co-occurrence distribution of the original mapping set containing pairs with n and S values lower than these thresholds shows no significant difference between the original and random mappings, and therefore is considered unreliable. While this value pair seems appropriate to be used as official thresholds, we selected n≥ 2 and S > 0.2 to make a safe choice and eliminate all the initial Domain2GO mappings other than those with high reliability. Following this analysis, the finalized Domain2GO mapping set contained 26,696 associations between 4,742 InterPro domains and 11,742 GO terms.

**Table 2.**
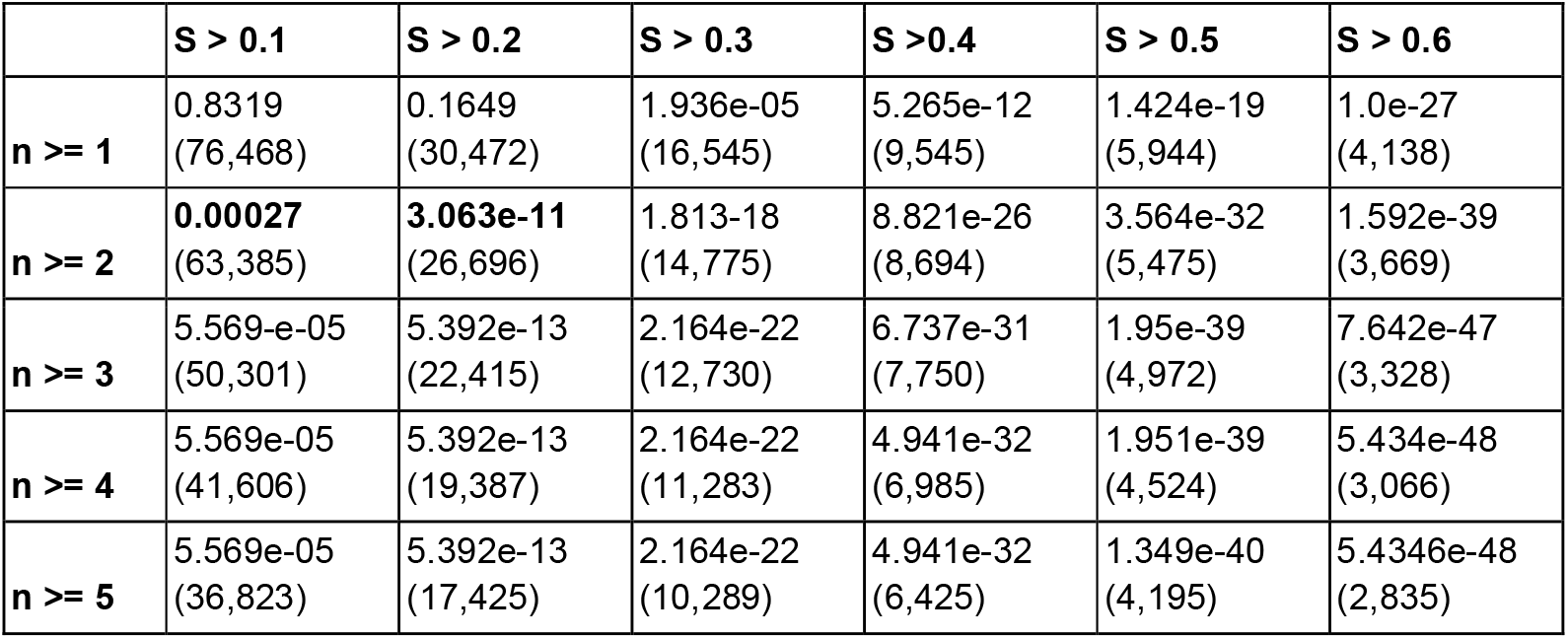
KS test significance results for the comparison of original vs. randomized co-occurrence similarity distributions at different co-occurrence similarity (S) and the number of co-annotated proteins (n) thresholds. The number of domain-GO mappings in each sample is given in parentheses.

|  | **S > 0.1** | **S > 0.2** | **S > 0.3** | **S > 0.4** | **S > 0.5** | **S > 0.6** |
| --- | --- | --- | --- | --- | --- | --- |
| **n >= 1** | 0.8319<br>(76,468) | 0.1649<br>(30,472) | 1.936e-05<br>(16,545) | 5.265e-12<br>(9,545) | 1.424e-19<br>(5,944) | 1.0e-27<br>(4,138) |
| **n >= 2** | **0.00027**<br>(63,385) | **3.063e-11**<br>(26,696) | 1.813-18<br>(14,775) | 8.821e-26<br>(8,694) | 3.564e-32<br>(5,475) | 1.592e-39<br>(3,669) |
| **n >= 3** | 5.569-e-05<br>(50,301) | 5.392e-13<br>(22,415) | 2.164e-22<br>(12,730) | 6.737e-31<br>(7,750) | 1.95e-39<br>(4,972) | 7.642e-47<br>(3,328) |
| **n >= 4** | 5.569e-05<br>(41,606) | 5.392e-13<br>(19,387) | 2.164e-22<br>(11,283) | 4.941e-32<br>(6,985) | 1.951e-39<br>(4,524) | 5.434e-48<br>(3,066) |
| **n >= 5** | 5.569e-05<br>(36,823) | 5.392e-13<br>(17,425) | 2.164e-22<br>(10,289) | 4.941e-32<br>(6,425) | 1.349e-40<br>(4,195) | 5.4346e-48<br>(2,835) |

### 3.2. Ablation study (enrichment analysis)

In order to assess the prediction capabilities of different versions of the domain-function mapping procedure, we employed three different measures calculated by the EM algorithm (as explained in Section 2.3) and compared their results to those of the co-occurrence similarity-based mapping. The first measure, which we denote as “θ_i_”, is an initial estimate of the probability of association between a domain and a GO term. The EM algorithm was performed to refine this initial estimate for each domain-GO term pair, and the output score is denoted as θ, and used as the second measure. The refined θ value is considered to be obtained at the point where the likelihood (Eq. 6) stops increasing noticeably. Finally, we performed a likelihood ratio test by rerunning EM for each pair, this time excluding the given pair from the pool of competing domain-GO term pairs (as explained in Section 2.3.2). The resulting likelihood ratio, which is denoted as E, stands as an evidence score for the given domain-GO term association, and is used as our third measure. To compare the prediction performance of each different statistical measure (i.e., S, θ_i_, θ, and E), ideally, the likelihood ratio test should have been performed for each Domain2GO pair in the initial set (2,069,796 mappings) by rerunning the EM until the likelihood values (Eq. 6) stop increasing noticeably. However, it was only possible to perform this step for a subset of the initial dataset composed of 78,477 mappings between 21,466 GO terms and 6,870 domains (obtained by filtering the initial set using a threshold of S ≥ 0.1) for 10 iterations due to high computational complexity (performing the same process for the full initial set would take 2,272 days on a single CPU of a regular consumer-level computer). Therefore, all the other measures were re-calculated based on this subset for comparison. We iterated EM algorithm 150 times to obtain the θ value, increasing the log-likelihood from −2.2 × 10^6^ to the stopping point of −5.89 × 10^5^. For the likelihood ratio test, we re-iterated EM algorithm 10 times for each domain-GO term pair, increasing the log-likelihood from −2.2 × 10^6^ to the stopping point of −6 × 10^5^ in each re-iteration.

An enrichment analysis was performed to evaluate which measure yields the most accurate associations. For this, we listed ranked predictions for each of the four measures (i.e., S, θi, θ and E). The number of ground truth domain-GO term pairs (provided by manually curated InterPro2GO associations) in the top K predictions was calculated for each measure. The number of manually associated pairs that are anticipated to appear in the top K predictions (i.e., expected mappings) was then estimated using the ratio of pairs that are verified by InterPro2GO across the entire prediction list. Enrichment plots (i.e., the ratio of observed to expected mappings in top K domain-GO term pairs) by each of the four measures (S, θi, θ and E) are displayed in Fig. 3. Pairs ranked by the θ_i_ scores showed no significant enrichment at any rank cutoff. The enrichment was 6.1-folds in the top 100 predictions when the pairs were ranked by θ, verifying the improved performance after EM optimization. Ranking by E increased the enrichment slightly further (from 3.4 to 4.7 fold) in the top 1000. The ranking by S yielded the highest enrichment, with 19 fold, in the top 100 predictions. The results suggest that the S score was able to identify significant domain-GO term pairs better than θ_i_, θ, and E, especially at low-rank cutoffs (in the top 100 to 10,000 predictions). However, if the likelihood ratio test could be performed on the full initial set with a sufficient number of iterations, it might be possible for the E score to compete with the S score. Based on the results of the ablation study, we decided to continue our experiments with the mappings obtained from co-occurrence similarity-based (S) filtering.

**Figure 3.**
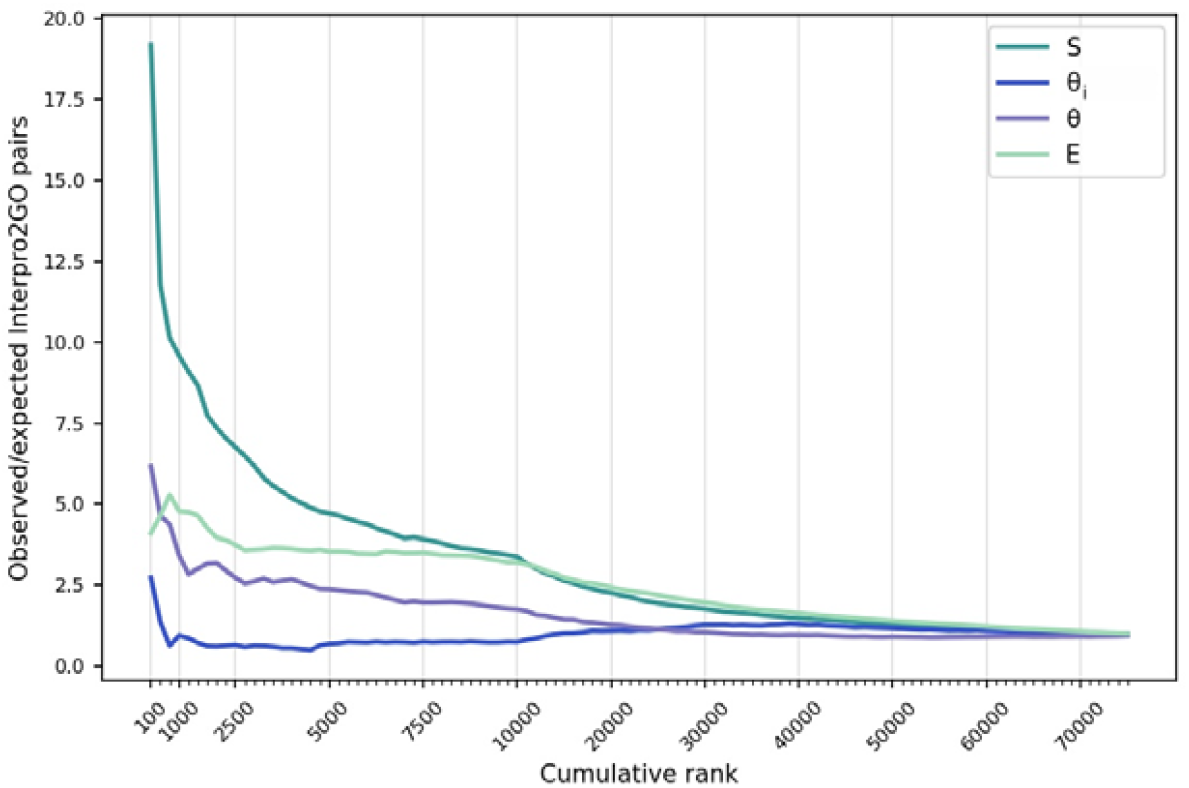
The enrichment of manually curated domain-GO associations in the top-ranking Domain2GO mappings. Mappings are ranked by four different measures: co-occurrence similarity (S), initial estimate of the association probability (θi), final estimate of the association probability (θ) and evidence measure for the association (E). The ratio of observed/expected manual associations in the top K mappings (y-axis) ranked by each of the four measures is plotted against the cumulative rank (x-axis). Higher enrichment values are indicators of better performance.

### 3.3. Comparison with the manually curated associations

The InterPro team has generated a set of curated domain function annotations called “InterPro2GO”, by first matching the InterPro entries with UniProtKB proteins using sequence information, and then assigning GO terms to them based on common features shared between the matched proteins (Camon et al., 2005). InterPro2GO mappings are manually curated, thus considered reliable. However, their coverage is low due to the time and labor-intensive nature of manual curation. In its current form, there are 5,881 manual associations between 3,056 domains and 1,524 GO terms in InterPro2GO (version: 2020/12).

To evaluate our method in terms of generating biologically relevant domain-GO associations, we compared the finalized Domain2GO mappings with InterPro2GO, as a reference domain annotation system. For this, we calculated the correspondence between domain-GO term pairs, which revealed that our predictions retrieved 737 out of 5,881 InterPro2GO associations. When the correspondence between domains was evaluated, we observed that 60% of the domains in the manual associations were paired with at least one GO term in our predictions as well. We found that 30% of the domains in our predictions were also included in the InterPro2GO associations. Similar calculations were performed on GO terms, yielding a 67% and 9% correspondence rate, respectively. According to these results, Domain2GO managed to find connections to more than half of the domains and GO terms presented in InterPro2GO, and many more.

Camon et al. followed a curation protocol in which they assigned GO terms to domains based solely on published experimental evidence of the domain’s specific function, thereby omitting numerous promising associations. As a result, the specificity of the protocol increased while its coverage decreased. Many of the associations retrieved by Domain2GO contain GO terms that are closely related (on the directed acyclic graph of Gene Ontology) to GO terms annotated to the same domain in InterPro2GO. For example, Interpro2GO mappings contain an association between “histone acetylation” (GO:0016573) and “nuclear receptor coactivator, CREB-bp-like, interlocking” (IPR014744), which was one of the most frequent domains in the manual associations. One of the GO terms we predicted to be associated with IPR014744 was “histone H2B acetylation” (GO:0043969), which is directly related (with “is a” relation) to the manually associated GO:0016573 as its child term. This comparison demonstrates that the proposed method was able to; (i) retrieve many of the mappings in InterPro2GO and (ii) provide more specific domain-function associations.

Table 3 displays 50 arbitrarily selected domain-GO term pairs with high reliability, based on their respective S and n values. As observed from the table, pairs with a high S score are commonly present in manually curated InterPro2GO mappings. When the correspondence between the manual associations and the high confidence mappings (i.e., pairs with an S ≥ 0.9) was calculated, it was found that 123 out of 737 high confidence mappings are found in InterPro2GO.

**Table 3.** Arbitrarily selected high-confidence domain-GO term pairs ranked by their S score. Pairs that are manually associated in the InterPro2GO set are marked with a “⍰”. *These pairs are included in Table 3 since they are examined as case studies in order to assess the biological relevance of mappings (see Section 3.4).

| Rank of the mapping | InterPro ID | GO ID | $\theta_i$ | $\theta$ | E | n | S | Are the domain & GO term manually associated? |
| --- | --- | --- | --- | --- | --- | --- | --- | --- |
| 1 | IPR000426 | GO:0019773 | 0.022 | 0.431 | 7.783 | 127 | 1 | ✓ |
| 2 | IPR039430 | GO:0046077 | 0.077 | 0.055 | 0.502 | 44 | 1 |  |
| 3 | IPR020602 | GO:0003934 | 0.019 | 0.171 | 1.448 | 40 | 1 |  |
| 4 | IPR000965 | GO:0004350 | 0.008 | 0.276 | 17.484 | 39 | 1 | ✓ |
| 5 | IPR011095 | GO:0008716 | 0.006 | 0.176 | 2.15 | 36 | 1 | ✓ |
| 6 | IPR003526 | GO:0008685 | 0.006 | 0.496 | 33.076 | 31 | 1 | ✓ |
| 7 | IPR032678 | GO:0006423 | 0.029 | 0.275 | 1.13 | 59 | 0.992 |  |
| 8 | IPR022630 | GO:0004478 | 0.143 | 0.208 | 0.69 | 57 | 0.991 | ✓ |
| 9 | IPR020598 | GO:0000179 | 0.028 | 0.401 | 3.215 | 56 | 0.991 | ✓ |
| 10 | IPR033828 | GO:0003883 | 0.031 | 0.067 | 0.807 | 52 | 0.99 | ✓ |
| 11 | IPR033705 | GO:0006438 | 0.077 | 0.253 | 1.515 | 50 | 0.99 |  |
| 12 | IPR039430 | GO:0046072 | 0.077 | 0.055 | 0.502 | 44 | 0.989 |  |
| 13 | IPR022417 | GO:0004418 | 0.017 | 0.294 | 3.332 | 43 | 0.989 | ✓ |
| 14 | IPR035490 | GO:0004360 | 0.008 | 0.087 | 1.301 | 42 | 0.988 |  |
| 15 | IPR033644 | GO:0004325 | 0.018 | 0.302 | 3.106 | 39 | 0.987 |  |
| 16 | IPR013840 | GO:0003911 | 0.03 | 0.068 | 0.771 | 38 | 0.987 | ✓ |
| 17 | IPR039429 | GO:0004372 | 0.008 | 0.059 | 1.123 | 69 | 0.986 |  |
| 18 | IPR013512 | GO:0051483 | 0.002 | 0.055 | 0.401 | 33 | 0.985 |  |
| 19 | IPR031481 | GO:0046920 | 0.003 | 0.394 | 4.066 | 33 | 0.985 |  |
| 20 | IPR018149 | GO:0004824 | 0.125 | 0.094 | 2.311 | 49 | 0.98 | ✓ |
| 21 | IPR033731 | GO:0004820 | 0.167 | 0.438 | 50.044 | 72 | 0.98 |  |
| 22 | IPR012310 | GO:0003910 | 0.031 | 0.129 | 2.273 | 46 | 0.979 | ✓ |
| 23 | IPR004971 | GO:0004482 | 0.182 | 0.403 | 7.345 | 21 | 0.977 |  |
| 24 | IPR020630 | GO:0004486 | 0.042 | 0.097 | 1.513 | 60 | 0.976 |  |
| 25 | IPR033659 | GO:0004325 | 0.018 | 0.195 | 1.093 | 38 | 0.974 |  |
| 26 | IPR011128 | GO:0047952 | 0.017 | 0.24 | 2.728 | 55 | 0.973 |  |
| 27 | IPR022663 | GO:0008839 | 0.016 | 0.154 | 0.779 | 33 | 0.971 | ✓ |
| 28 | IPR041739 | GO:0004349 | 0.008 | 0.231 | 6.538 | 31 | 0.969 |  |
| 29 | IPR004255 | GO:0004144 | 0.016 | 0.097 | 0.436 | 15 | 0.968 | ✓ |
| 30 | IPR005110 | GO:0042040 | 0.003 | 0.081 | 0.266 | 15 | 0.968 |  |
| 31 | IPR001442 | GO:0005587 | 0.022 | 0.149 | 0.612 | 15 | 0.968 |  |
| 32 | IPR005110 | GO:0018315 | 0.045 | 0.081 | 0.266 | 15 | 0.968 |  |
| 33 | IPR007471 | GO:0004057 | 0.167 | 0.18 | 0.561 | 15 | 0.968 | ✓ |
| 34 | IPR039430 | GO:0009189 | 0.015 | 0.055 | 0.502 | 44 | 0.967 |  |
| 35 | IPR015341 | GO:0004559 | 0.016 | 0.151 | 1.435 | 29 | 0.967 | ✓ |
| 36 | IPR035490 | GO:0070548 | 0.033 | 0.087 | 1.301 | 42 | 0.966 |  |
| 37 | IPR011258 | GO:0004619 | 0.005 | 0.242 | 1.469 | 28 | 0.966 | ✓ |
| 38 | IPR009721 | GO:0047196 | 0.083 | 0.089 | 0.346 | 14 | 0.966 |  |
| 39 | IPR022673 | GO:0004340 | 0.008 | 0.065 | 0.611 | 41 | 0.965 |  |
| 40 | IPR020602 | GO:0006729 | 0.004 | 0.171 | 1.448 | 40 | 0.964 |  |
| 41 | IPR041711 | GO:0019988 | 0.023 | 0.093 | 0.914 | 39 | 0.963 |  |
| 42 | IPR005849 | GO:0008108 | 0.027 | 0.21 | 0.62 | 13 | 0.963 | ✓ |
| 43 | IPR018164 | GO:0004813 | 0.029 | 0.066 | 1.284 | 63 | 0.962 | ✓ |
| 44 | IPR022648 | GO:0044796 | 0.007 | 0.127 | 0.72 | 25 | 0.962 |  |
| 45 | IPR035476 | GO:0004347 | 0.008 | 0.118 | 1.813 | 49 | 0.961 |  |
| 46 | IPR013548 | GO:0002116 | 0.006 | 0.051 | 0.277 | 24 | 0.96 |  |
| 47 | IPR000070 | GO:0030599 | 0.005 | 0.345 | 23.678 | 69 | 0.958 | ✓ |
| 48* | IPR001017 | GO:0004591 | 0.005 | 0.035 | 0.325 | 22 | 0.557 |  |
| 49* | IPR001678 | GO:0008169 | 0.03 | 0.017 | 0.164 | 24 | 0.387 |  |
| 50* | IPR003583 | GO:0051103 | 0.25 | 0.015 | 0.166 | 25 | 0.394 |  |

### 3.4. Biological relevance of the selected mappings—a case study

To evaluate the biological significance of our findings, we examined three Domain2GO mappings. The statistics of these domain-GO pairs are shown in the last 3 rows of Table 3. Also, the semantic relationships between predicted and manually annotated GO terms for each domain entry are shown for each case in different panels of Fig. 4.

**Figure 4.**
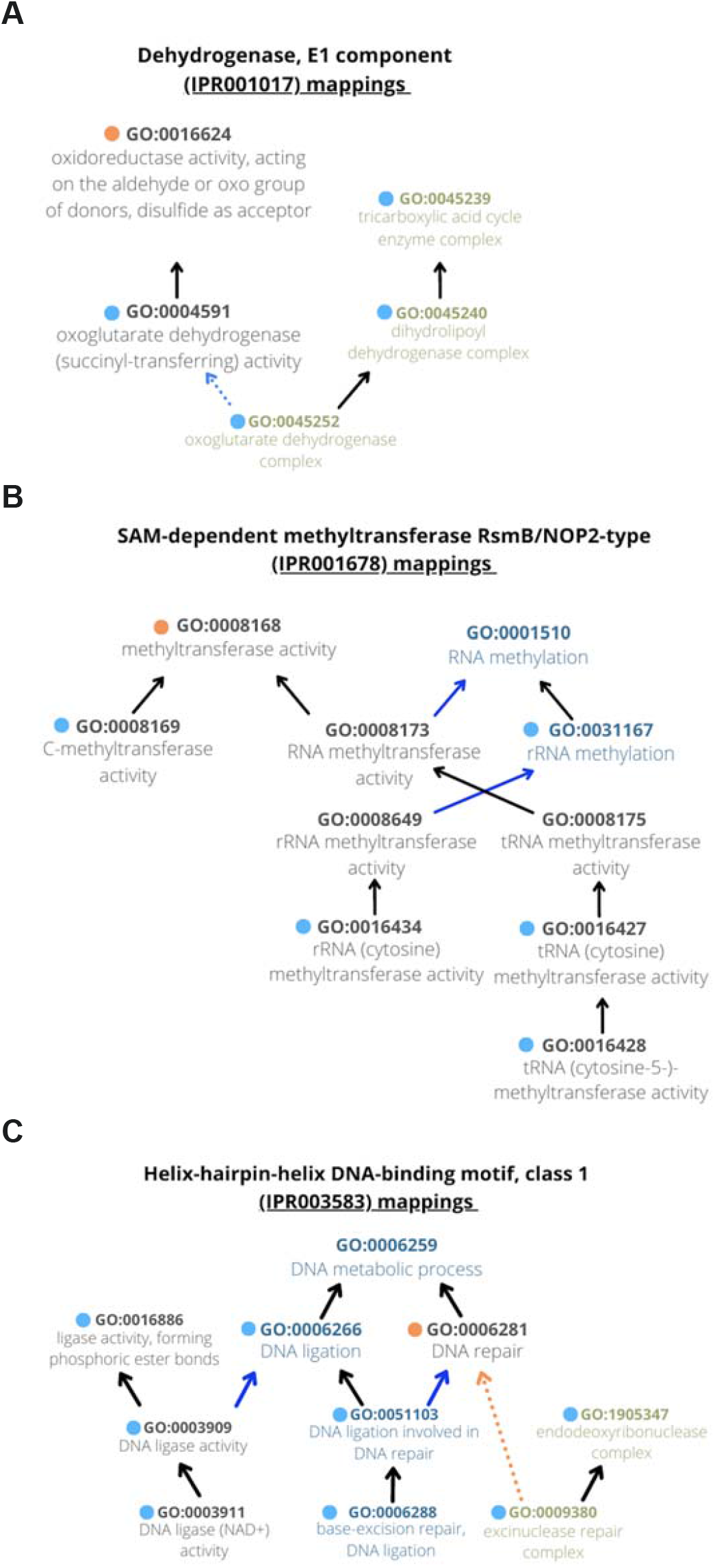
A case study by comparing Domain2GO with curated InterPro2GO annotations on selected domains. **(A)** GO terms associated with Dehydrogenase, E1 component (IPR001017) domain. **(B)** GO terms associated with SAM-dependent methyltransferase RsmB/NOP2-type (IPR001678) domain. **(C)** GO terms associated with Helix-hairpin-helix DNA-binding motif, class 1 (IPR003583) domain.

The first case is a Domain2GO mapping between the domain “dehydrogenase, E1 component” (IPR001017) and the GO molecular function term “oxoglutarate dehydrogenase (succinyl-transferring) activity” (GO:0004591). These two terms were mapped to each other with a co-occurrence similarity (S) value of 0.56, on a total of 22 proteins (n). According to the InterPro database, IPR001017 domain is found in multiple proteins that have dehydrogenase activity, one of which is the 2-oxoglutarate dehydrogenase (EC 1.2.4.2). (URL:https://www.ebi.ac.uk/interpro/entry/InterPro/IPR001017/) The E1 subunit catalyzes the decarboxylation of this enzyme by transferring a succinyl group onto the lipoyl-group bound to the E2 subunit (Tretter & Adam-Vizi, 2005). Clearly, this domain is in charge of the molecular function defined by the Domain2GO mapped GO term. There is an InterPro2GO generated association between IPR001017 and the molecular function term “oxidoreductase activity, acting on the aldehyde or oxo group of donors, disulfide as acceptor” (GO:0016624), which is related (i.e., with is_a relationship) to the Domain2GO predicted GO:0004591 as its parent term. Additionally, Domain2GO captured another association between IPR001017 and the cellular component term “oxoglutarate dehydrogenase complex” (GO:0045252), which is a multienzyme complex that contains 2-oxoglutarate dehydrogenase as one of its main components (Bunik & Degtyarev, 2008). GO:0045252 is also related to GO:0004591 (i.e., with capable_of relationship). This example shows that Domain2GO was able to associate IPR001017 with GO terms that are more specific compared to the manually curated GO association. In addition to that, we found additional associations between this domain and different cellular component terms, each of which is one-step away from the above-mentioned terms (and from each other) on the GO DAG (Fig. 4A).

The second example case is a mapping between the domain “SAM-dependent methyltransferase RsmB/NOP2-type” (IPR001678) and the molecular function term “C-methyltransferase activity” (GO:0008169). These two terms were mapped to each other with a co-occurrence similarity (S) value of 0.39, on a total of 24 proteins (n). The “methyltransferase activity” of the IPR001678 domain is verified by a manually curated InterPro2GO association with the GO term of the same name (GO:0008168). GO:0008168 is also related (i.e., with is_a relationship) to the Domain2GO mapped GO:0008169 as its direct parent. As the names and the parent-child relationship of the two terms suggest, “C-methyltransferase activity” defines a more specific function. The InterPro database description of the IPR001678 domain is as follows: “The C-terminal domain of ribosomal RNA cysteine methyltransferases is highly conserved in archaeal, bacterial, and eukaryotic proteins” (URL: https://www.ebi.ac.uk/interpro/entry/InterPro/IPR001678/). The proteins that are mentioned in this description could be associated with the “C-methyltransferase activity” function thanks to the IPR001678-GO:0008169 mapping in Domain2GO. Also, as shown in Fig. 4B, we were able to find links between IPR001678 and some other molecular function and biological process terms that are also closely related to the InterPro2GO associated GO:0008168 term.

The last example case is a mapping between the domain “helix-hairpin-helix DNA-binding motif, class 1” (IPR003583) and the GO biological process term “DNA ligation involved in DNA repair” (GO:0051103). These two terms were mapped to each other with S=0.39 and n=25. Helix-hairpin-Helix (HhH) is known as the most common nucleic-acid binding module in repair proteins (Aravind et al., 1999). The role of HhH domain in repair processes is also verified by a manually curated InterPro2GO association between IPR003583 and the biological process term “DNA repair” (GO:0006281), which is related (i.e., with part_of relationship) to the mapped GO:0051103 term as its parent. GO:0051103 is defined as a process that “contributes to DNA repair, carried out by DNA ligase” in the UniProt-GOA database (URL: https://www.ebi.ac.uk/QuickGO/term/GO:0051103). HhH was detected in DNA ligases at both sequence and structural levels by the elucidation of crystal structures (Doherty & Suh, 2000). Accordingly, the proteins that contain the domain IPR003583, and thus annotated to the function term GO:0051103 thanks to the corresponding Domain2GO mapping, are all DNA ligases from different organisms according to the information provided in UniProtKB (e.g., UniProt accessions: P15042, Q9WXV5, Q8A9C1, Q8DPS9, etc.). We also found a mapping between IPR003583 and the GO cellular component term “excinuclease repair complex” (GO:0009380), which is related (i.e., with capable_of_part_of relationship) to the manually associated GO:0006281 term, as its child. This is another clear example of Domain2GO associating domains with biologically relevant GO terms. Importantly, whereas InterPro2GO defines associations between the IPR003583 domain and two GO terms, one from the biological process and the other from the molecular function categories, the proposed method was able to define additional associations with cellular component terms (Fig. 4C).

### 3.5. Protein function prediction performance evaluation

The main aim of Domain2GO is to predict the functions of domains, not full proteins. Nevertheless, it is interesting to examine the capabilities of Domain2GO in this regard. The full protein function prediction performance of Domain2GO was evaluated using the CAFA3 challenge benchmark dataset (i.e., proteins that have gained manually curated GO associations during the annotation collection period), and compared against 3 different methods (i.e., two CAFA baselines, BLAST and Naive classifier, and the manually curated InterPro2GO annotations). We tested two individual sets of protein function predictions for Domain2GO. The first one, “Domain2GO-S”, was generated via propagating the domain-GO mappings that were filtered with an S-score ≥ 0.05, to the proteins in the CAFA benchmark dataset. We assigned the co-occurrence similarity score of the respective domain-GO pair as the probabilistic prediction score of the corresponding protein-GO association prediction. By filtering mappings based on S-score ≥ 0.05, we eliminated non-reliable domain-GO mappings and still ended up with a prediction set of reasonable coverage (resulting in 169,534 Domain2GO mappings, which led to 8,695,580 function predictions between 24,665 GO terms and 75,354 CAFA3 target proteins). For the second set, “Domain2GO-E”, we generated protein-function predictions this time using Domain2GO mappings previously used as the input of the EM algorithm (78,477 mappings, as explained in section 3.2) and assigned their E-value (i.e., the output of the EM algorithm) as the probabilistic scores of the corresponding protein-GO association predictions. This set contained 2,989,529 predictions between 21,187 GO terms and 73,860 target proteins. We evaluated the results based on the “no-knowledge benchmarks”, which contain proteins that did not have any experimentally curated annotations in any of the GO ontologies before the benchmark collection period.

We were unable to generate predictions for all target proteins due to the thresholding operations we performed beforehand on domain-GO mappings, and the lack of domain annotation for a part of the dataset. This loss impacted the performance of the proposed method in terms of coverage. The coverage of the Domain2GO-S, Domain2GO-E and InterPro2GO on the CAFA3 no-knowledge benchmark set was %0.53, %0.43 and %0.22, respectively (i.e., C in Fig. 5). Since Domain2GO predicts protein functions based only on domain information, it was expected that both Domain2GO-S and Domain2GO-E would get lower coverage values than the baseline methods. InterPro2GO had the lowest coverage in each GO ontology, which makes sense since it only made predictions based on experimental evidence.

**Figure 5.**
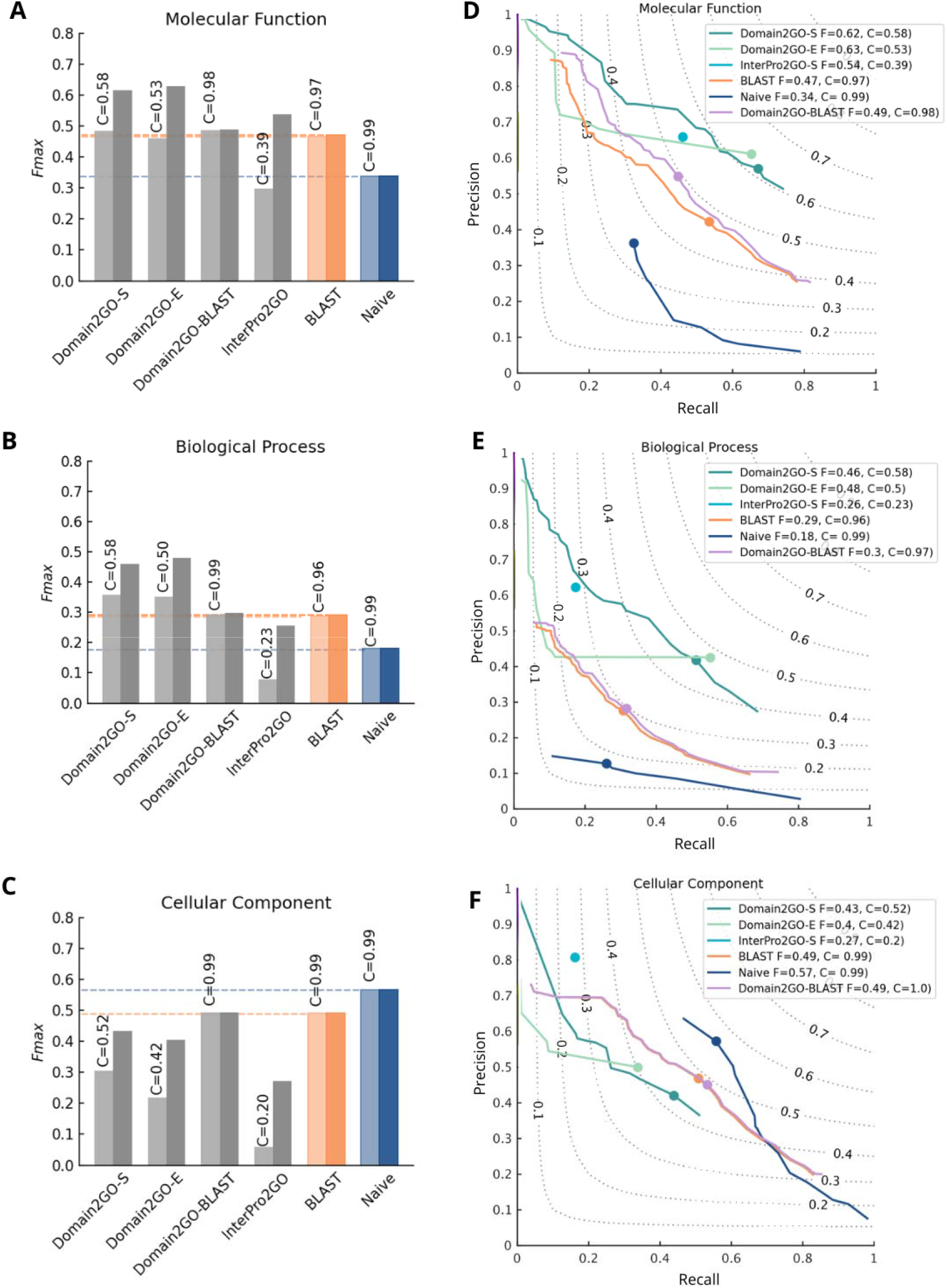
Protein function prediction performance results (Fmax) on the CAFA3 benchmark set for 3 ontologies. A-C: Fmax bar plots for all methods. The light-colored bars represent the performance in the full mode, while the dark-colored bars represent the performance in the partial mode. The coverage of each method is shown as text on top of the bars. **D-F: Precision-recall curves for all methods in the partial evaluation mode**. The dot on the curve is where the maximum Fmax score is reached. The perfect predictor should achieve Fmax=1, which would be located at the top right corner of the plot.

Fig. 5 shows the performance comparison of Domain2GO with InterPro2GO and baseline methods, in terms of *Fmax* values (over the entire CAFA3 benchmark dataset, i.e., the full mode). Both Domain2GO-S and Domain2GO-E achieved better performances compared to the InterPro2GO and baseline predictors when predicting molecular function and biological process terms (*Fmax* = 0.48 and 0.46 for MFO and *Fmax* = 0.36 and 0.35 for BPO, respectively). In the case of predicting cellular component terms, Domain2GO-S, Domain2GO-E and InterPro2GO models performed worse compared to the baseline methods, with a performance of *Fmax* = 0.31, 0.22 and 0.06 respectively. To eliminate the impact of the low coverage on performance scores, we also analyzed the predictions in partial mode. In this evaluation mode, prediction methods are only judged on the proteins for which they make predictions. The Fmax values and the precision-recall curves of the evaluation in the partial mode are shown in Fig. 5D and E. The performance of all three models with low coverage increased significantly when the predictions were evaluated in the partial mode. An interesting observation here is that the baseline methods still outperform the domain-based predictors in the CCO category in partial mode. A few possible reasons for obtaining different performances among the three ontologies could be: (i) differences in the structure of the three ontologies and their indicator data types; (ii) the annotation status (completeness) of benchmark proteins; and (iii) the biases in the nature of manually curated protein function annotations (Peng et al., 2018). The most likely reason behind the relatively low predictive performance of Domain2GO and InterPro2GO in the CCO category could be that it is harder to associate domains with cellular structures, as a domain can be part of various proteins functioning in different locations.

In general, baseline methods are prone to increasing the number of true positives by correctly predicting a small number of relatively generic (shallow) terms, which can be the reason for the competitive performance of baselines (especially the Naive classifier) in the CCO category under the Fmax evaluation (Jiang et al., 2016). Smin scores can provide insight into the capability of a model to make accurate but also specific predictions, as this measure depends on the information content of predicted GO terms. As shown in Fig. 6, the exceptionally high Fmax performance of the Naive classifier in the CCO category is degraded once the relatively general terms are down-weighted in the Smin-based evaluation (lower Smin values indicate higher performance). In partial evaluation mode, Domain2GO-S achieved a better performance compared to baseline predictors, with an Smin of 5.62, 15.34, and 5.53 for MFO, BPO, and CCO, respectively (Fig. 6). InterPro2GO was able to accurately predict highly specific CCO terms as it extracted GO terms from curated annotations obtained from articles, which resulted in a better performance compared to Domain2GO-S with an Smin of 5.14, 15.13, and 4.87 for MFO, BPO, and CCO, respectively.

**Figure 6.**
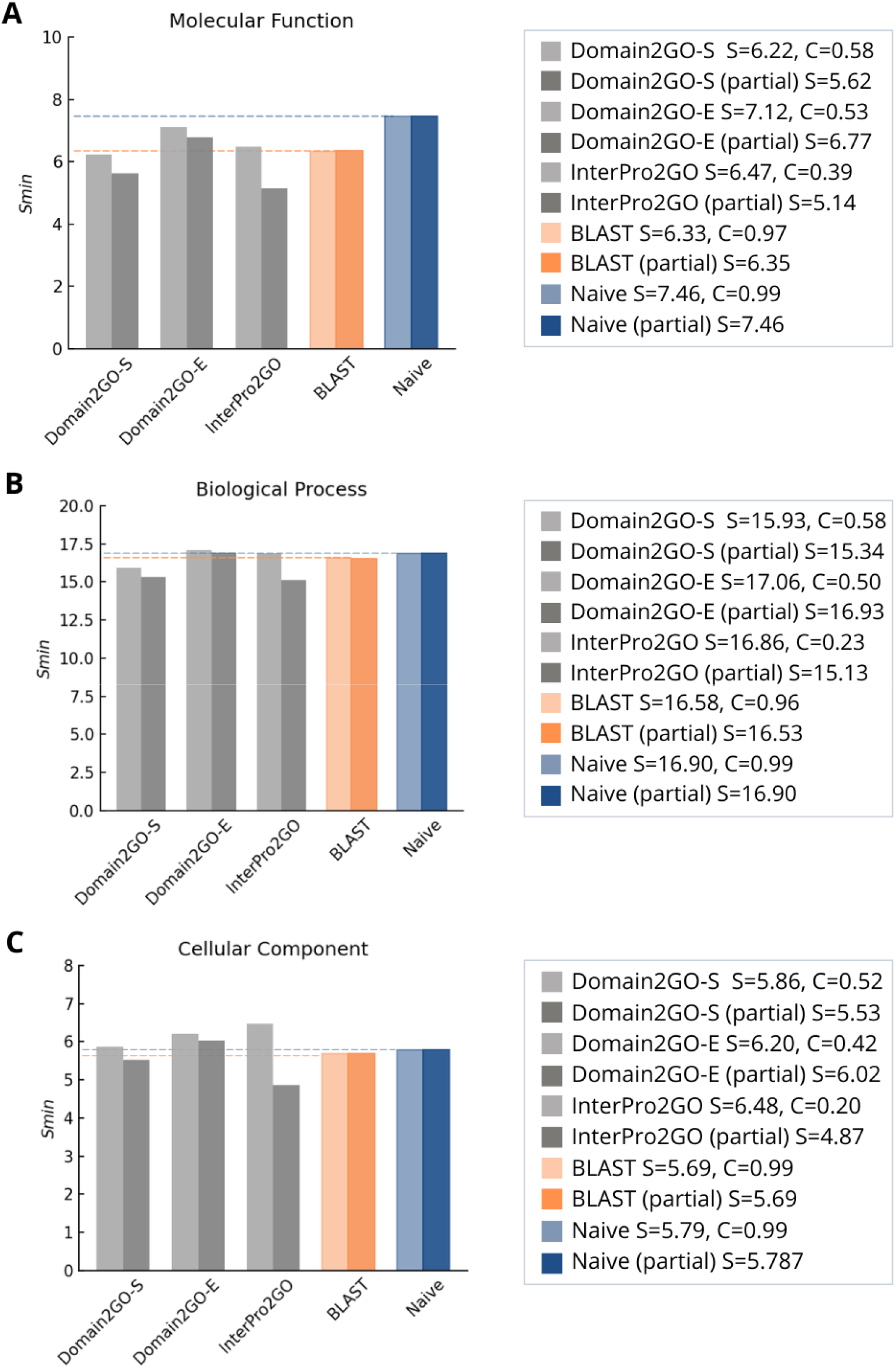
Protein function prediction performance results (Smin) on the CAFA3 benchmark set for 3 ontologies. **(A)** molecular function ontology, **(B)** biological process ontology, and **(C)** cellular component. The light-colored bars represent the performance in the full mode, while the dark-colored bars represent the performance in the partial mode. The coverage of each method is shown as text on top of the bars. Lower minimum semantic distance (Smin) values indicate higher performance.

We created a hybrid protein function prediction approach that utilizes both sequence and domain-based data, with the aim of investigating the correspondence between these two types of approaches and the impact of combining them. For this, a consensus of Domain2GO-S and BLAST prediction sets was deduced. The so called “Domain2GO-BLAST” prediction set was made up of every protein-GO term pair that was predicted by either the Domain2GO-S method or the BLAST method, or by both. When BLAST-only and Domain2GO-only predictions are considered, the Domain2GO-BLAST prediction scores are the same as the scores from the prediction set from which this pair was taken. For the pairs contained in both Domain2GO and BLAST prediction sets, if a pair is predicted as negative by both Domain2GO-S and BLAST at a certain threshold, the Domain2GO-BLAST prediction for this pair is assigned as negative at this threshold. In all other situations, the Domain2GO-BLAST prediction for this pair is assigned as positive at this threshold. The CAFA3 benchmark analysis on CCO using Domain2GO-BLAST predictions (i.e., *Fmax* = 0.493) resulted in a significant improvement over Domain2GO-S and a slightly better performance compared to BLAST (Fig. 5). The performances for MFO and BPO (i.e., *Fmax* = 0.49 and 0.30, respectively) are similar to those of the original Domain2GO-S and BLAST (Fig. 5).

Finally, we comparatively examined Domain2GO and BLAST in terms of their intersecting and differing true-positive, false-positive, and false-negative predictions at the optimal thresholds (i.e., the threshold where Fmax is obtained at) for all three ontologies (Fig. 7). Even though Domain2GO’s low coverage meant that it made fewer true-positive predictions than BLAST, it also made a lot fewer false-positive predictions. This gave Domain2GO-S a better overall performance and shows how precise this method is. The high number of mutual predictions in all three ontologies shows that Domain2GO-S and BLAST are similar, which explains why there aren’t significant differences in performance when comparing the Domain2GO-BLAST prediction set to Domain2GO-S and BLAST on their own. The performance of Domain2GO-BLAST in BPO, which was slightly better than the performance of BLAST but much lower than the performance of Domain2GO-S, can be explained by the large number of false-positive predictions made by BLAST alone.

**Figure 7.**
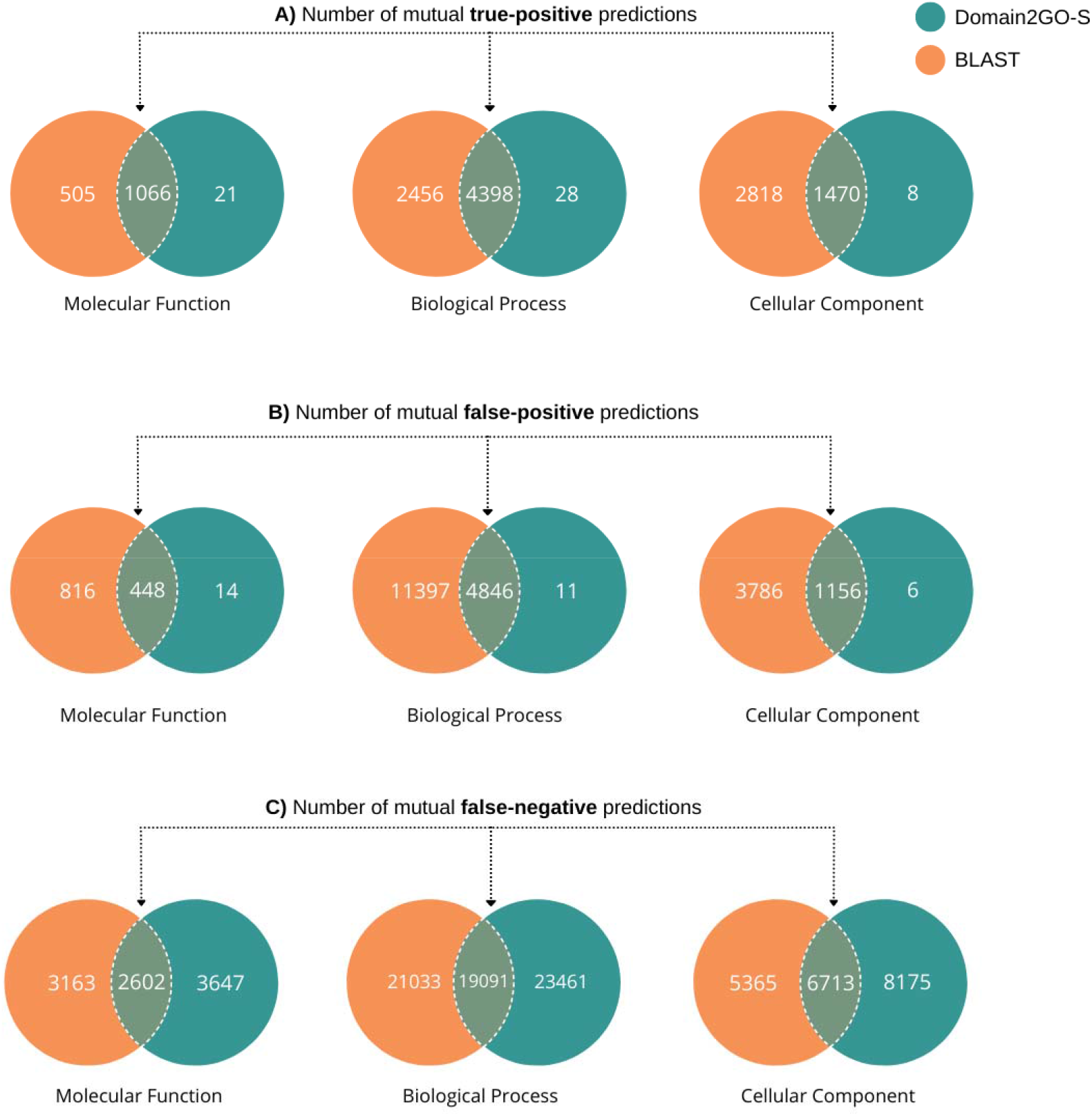
The analysis of correspondence between Domain2GO and BLAST predictions. The number of: (A) true positives, (B) false positives, (C) false negatives of Domain2GO-S-only, mutual, and BLAST-only predictions at the optimal thresholds. The mutual positive predictions are the protein-GO term pairs, which are predicted as positive by either the Domain2GO-S or BLAST methods. The mutual negative predictions are the protein-GO term pairs, which are predicted as negative by both the Domain2GO-S and BLAST methods.

### 3.6. Generation of the finalized protein function predictions

Finally, we employed final Domain2GO mappings to predict GO term annotations for 566,996 proteins in the UniProtKB/SwissProt database (release 2022_01), by propagating domain-associated GO terms to the proteins that contain those domains, based on annotations provided in the InterPro database. The resulting protein function prediction set contains 5,046,060 GO term predictions for 291,519 proteins and 11,742 GO terms, which are available at https://github.com/HUBioDataLab/Domain2GO.

## 4. Conclusion

In this study, we proposed Domain2GO; a straightforward domain function prediction approach that maps structural domains of proteins to function-defining GO terms. Two main statistical measures; co-occurrence similarity and E value, were calculated in order to quantitatively define the association patterns of domains and functions in proteins. An enrichment analysis-based ablation study was performed on manually curated domain function annotations to assess the accuracy of different statistical measures utilized in Domain2GO. Additionally, a literature-based case study was performed to investigate the significance of Domain2GO mappings.

Even though Domain2GO is not specifically designed to predict protein functions, we looked at how well it did in this area. For this, the domain-GO mappings were utilized to generate protein function predictions, which were then evaluated on the CAFA3 challenge benchmark set. The results revealed that Domain2GO outperforms InterPro2GO; a domain-function mapping approach based on experimental evidence, as well as two baseline methods presented in CAFA challenges (in terms of predicting terms in the biological process and molecular function ontologies), which can be considered successful considering that the main aim of Domain2GO is not protein function prediction. Furthermore, a simple hybrid function prediction approach, which combines domain-based Domain2GO and sequence-based BLAST, performed better than Domain2GO on the cellular component ontology and BLAST on the molecular function ontology, indicating the benefit yielded from combining different data sources and exploiting a larger fraction of the knowledge space.

It is possible to produce large-scale GO annotations for unknown proteins using the domain-GO mappings produced in this study. This process can be easily scaled up to the entire UniProtKB (i.e., nearly 227 million protein entries), which is a critical issue for most of the state-of-the-art protein function prediction methods. For example, in this study, the prediction of GO annotations for the UniProtKB/Swiss-Prot database (i.e., 291,519 proteins in it) only took 4 minutes on a regular consumer-level computer.

In the field of predictive modeling of biology, using a single type of data is often limited by the fact that that data can only tell you about one aspect of the prediction task at hand. Thus, the limitations we encountered in this regard (e.g., while predicting cellular component terms) can be overcome by incorporating different data sources that provide insight into the functional aspects that are unpredictable by the domain-based approach. In this sense, a potential area of exploration could be extending the Domain2GO framework from merely using domain-based data to include various complementary data sources. Furthermore, the biological entity association approach proposed in this study can be modified with the use of additional types of features and ontologies in order to develop novel multi-modal systems that can predict unknown biomolecular relationships.

It is possible to utilize Domain2GO via the open access data repository of the project, which contains all datasets, results, and the source code at https://github.com/HUBioDataLab/Domain2GO.

